# Visualization and quantification of spatiotemporal disease progression in Arabidopsis using a bioluminescence-based imaging system

**DOI:** 10.1101/2024.10.09.617450

**Authors:** Nanne W. Taks, Mathijs Batstra, Ronald Kortekaas, Floris D. Stevens, Sebastian Pfeilmeier, Harrold A. van den Burg

## Abstract

Plant pathogenic bacteria use various entry strategies to colonize their host, like entering through natural openings and wounds in leaves and roots. The vascular pathogen *Xanthomonas campestris* pv. campestris (Xcc) enters through hydathodes, organs at the leaf margin involved in guttation. Subsequently, Xcc breaks out from infected hydathodes, progressing into the xylem vessels and causing systemic disease. To elucidate the mechanisms that underpin the different stages of Xcc pathogenesis, a need exists to image Xcc progression *in planta* in a non-invasive manner. Here, we describe a phenotyping setup and Python image analysis pipeline capturing the Xcc infection in 16 *Arabidopsis thaliana* plants in parallel over time. The setup used both an RGB to capture disease symptoms and an ultra-sensitive CCD camera to monitor bacterial progression inside the leaves using bioluminescence. We demonstrate that the image analysis pipeline reliably quantifies bacterial growth *in planta* for two bacterial species, that is vascular Xcc and the mesophyll pathogen *Pseudomonas syringae* pv. tomato. The resolution of the camera allowed early detection of Xcc in the hydathodes, yielding valuable information on this early stage of the Xcc infection process. The data obtained through the automated image analysis pipeline was robust and validated findings from other bioluminescence imaging methods, while requiring fewer samples. We can thus quantify the resistance level of a large number of *Arabidopsis thaliana* accessions and mutant lines to different bacterial strains in a non-invasive manner for phenotypic screenings.

## Introduction

Plants are colonized by a wide range of pathogenic microbes that apply different strategies to infect plant tissues (Jian et al., 2024). To study these interactions, it is important to reliably quantify plant susceptibility to a specific pathogen. For phytopathogenic bacteria, such as the model pathogen *Pseudomonas syringae* pv. tomato (Pst), standard phenotyping methods still rely on visual assessment of plant responses and the quantification of bacterial colonization via colony-forming units (CFUs)(Tornero and Dangl, 2001; Whalen et al., 1991). While these methods typically provide accurate evaluations of the level of plant disease resistance and susceptibility, they are often labor-intensive and destructive. This poses challenges when studying pathogens that follow a multi-staged infection process accross different tissues, as these methods often disregard spatial interactions over time. A prime example can be found in the vascular bacterial pathogen *Xanthomonas campestris* pv. campestris (Xcc), an economically relevant pathogen of cabbage that shows several distinct stages during its infection cycle (Cerutti et al., 2017; Paauw et al., 2023). This specialized bacterium exploits hydathodes, which are gland-like structures at the leaf margin that excrete xylem sap in a process called guttation, as its initial entry point into plant tissue (Cerutti et al., 2017; Hugouvieux et al., 1998). Inside the hydathode, the bacteria are found to proliferate for up to a week (depending on the level of plant resistance) without causing any visual disease symptoms (Paauw et al., 2023). They then transit into the xylem vessels that protrude into the hydathode tissue and systemically spread from these points to the rest of the vasculature and across the leaf (Cerutti et al., 2017; Paauw et al., 2023). Earlier work has shown that *Arabidopsis thaliana* (hereafter Arabidopsis) displays an immune response both in the hydathodes and vasculature upon Xcc recognition, which restricts bacterial spread at different stages of infection (Lin et al., 2022; Paauw et al., 2023; Taks et al., 2024; Wang et al., 2015). To study the mechanism underpinning Xcc colonization of different tissues in Arabidopsis, a need exists to develop imaging methods that allow us to visualize the entire infection process in one individual plant over time in a non-invasive manner.

Currently, methods to quantify the spread and proliferation of Xcc in Arabidopsis include manual scoring of disease symptoms using indices (Guy et al., 2013; Meyer et al., 2005; Wang et al., 2015), inferring the bacterial titer from sections of leaf tissue (Feng et al., 2012; Guy et al., 2013; Lin et al., 2022; Wang et al., 2015) and measuring the length of the V- shaped chlorotic lesions (Xie et al., 2023), a typical disease symptom caused by Xcc. Although these methods provide sufficient accuracy, they disregard hydathode colonization as a critical initial stage of the infection. In recent years, efforts have been made to visualize Xcc in hydathodes, focusing mainly on using bacterial reporter systems such as bioluminescence. Application of an engineered *luxCDABE* operon from *Photorabdus luminescens* (Winson et al., 1998) has proven successful in visualizing Xcc in Arabidopsis hydathodes (Cerutti et al., 2017; Paauw et al., 2024, 2023; van Hulten et al., 2019). However, quantification of hydathode colonization is still labor-intensive as the sampling methods remained destructive (single leaf detachment). Hence, valuable data on the ability of the bacteria to spread from hydathode to the vascular system cannot be captured by repeated measures on the same plant. Therefore, non-invasive methods for visualizing early hydathode infections and subsequent colonization of leaf tissue are needed.

Non-invasive visualization of luminescence is typically done using a highly sensitive charge-coupled device (CCD) camera. While often used for detection of horseradish peroxidase (HRP) activity in immunoblots, it has also been successfully applied to detect a range of pathogenic bacteria in plants (Furci et al., 2021; Jutras et al., 2021; Mutka et al., 2016; Soldan et al., 2021; Wang et al., 2024; Xu et al., 2022). Although many laboratories are equipped with CCD camera setups, they are not suited to process large plant populations due to size constraints of the device. However, to be able to fully exploit the genetic diversity and resources that are available for Arabidopsis (Simon et al., 2008; Van de Weyer et al., 2019; Weigel and Mott, 2009), high-throughput devices are needed to effectively screen large groups of plants. For instance, the resistance gene *SUT1* that acts in hydathodes against infection by Xcc (Taks et al., 2024) was identified by screening thousands of individual plants and evaluating the level of resistance by manual scoring of a luminescence index (van Hulten et al., 2019). To surpass such labor-intensive and bias prone methods, a bioluminescence imaging setup that enables screening of a plant population would significantly speed up these processes.

Quantification of bioluminescence is often performed using manual annotation of the images — either in proprietary software (Furci et al., 2021; Xu et al., 2022) or ImageJ (Jutras et al., 2021; Mutka et al., 2016; Soldan et al., 2021; Wang et al., 2024)— or with a routine image analysis pipeline. To avoid manual errors/bias, automated image analysis pipelines have been developed in ImageJ (Laflamme et al., 2016) and the OpenCV image analysis library in Python (Paauw et al., 2024). These pipelines offer fast quantification of disease symptoms or bacterial luminescence but require highly standardized input images. To acquire such images, plants need to be imaged in a rigid setup to allow for smooth image analysis processing. Therefore, a tailored system is required to be able to combine the power of bioluminescence and automated image analysis processing.

Here, to develop a system that enables both the tracking of the entire Xcc infection process in Arabidopsis and screening of large plant populations, a custom-made imaging cabinet was designed. This cabinet contains a dual camera, RGB and CCD, and allows for the successive imaging of the plants and the bioluminescence emitted by the bacteria inside these plants. In addition, we designed imaging trays that can hold 16 Arabidopsis rosettes in a grid allowing fully automated image processing. Finally, we developed a Python analysis pipeline to quantify both disease symptoms and bacterial spread. Here, we benchmark this method for Xcc and show that it can also be applied for other phytopathogenic bacteria. Our method holds promise for uncovering intricate mechanisms that happen in a highly localized and temporal basis during plant-pathogen interactions.

## Results

### How to build a digital phenotyper: designing the hardware

To assess disease severity of Xcc infection, we spray-inoculated Arabidopsis plants and applied humidity/temperature cycles to promote guttation and uptake of Xcc in hydathodes (van Hulten et al., 2019). Using a bioluminescent Xcc reporter strain, we then followed bacterial spread *in planta* for up to 14 days post inoculation (dpi) (Figure 1A). To unravel the complex interactions involving different host tissue types, i.e. hydathodes, xylem vessels and mesophyll, we aimed for a non-invasive method for imaging bacterial infection in a spatiotemporal manner. To this end, a custom-made phenotyping cabinet was designed that features white LEDs and two digital cameras, that is, an RGB camera for rosette imaging and an ultra-sensitive CCD camera for bioluminescence (Figure S1A). To avoid internal light reflections, the cabinet interior was painted black. As overlap between the plant rosettes can interfere with the automated image processing and analysis (Laflamme et al., 2016; Paauw et al., 2024), we designed a custom-made 4×4 tray using black anodized aluminum, which allows for the imaging of 16 plants in parallel (Figure S1B). In the case of Xcc infections, this ensures non-overlapping rosettes until at least 14 dpi. For other applications, in which smaller rosettes are imaged, trays with 40 or more rosettes can fit into the imaging cabinet. Plants can be directly grown in these trays or transferred prior to imaging. In the phenotyping cabinet, plants are imaged by an RGB camera (artificial lighting, 70 milliseconds exposure) and a CCD camera (darkness, 300 seconds exposure), resulting in two image files capturing the rosette phenotype and the spread of bacterial luminescence, respectively (Figure S1C). These images are then fed into our Python image analysis pipeline to obtain data on the individual plant sizes, disease symptoms (chlorosis) per plant and bacterial spread per plant.

**Figure 1.**
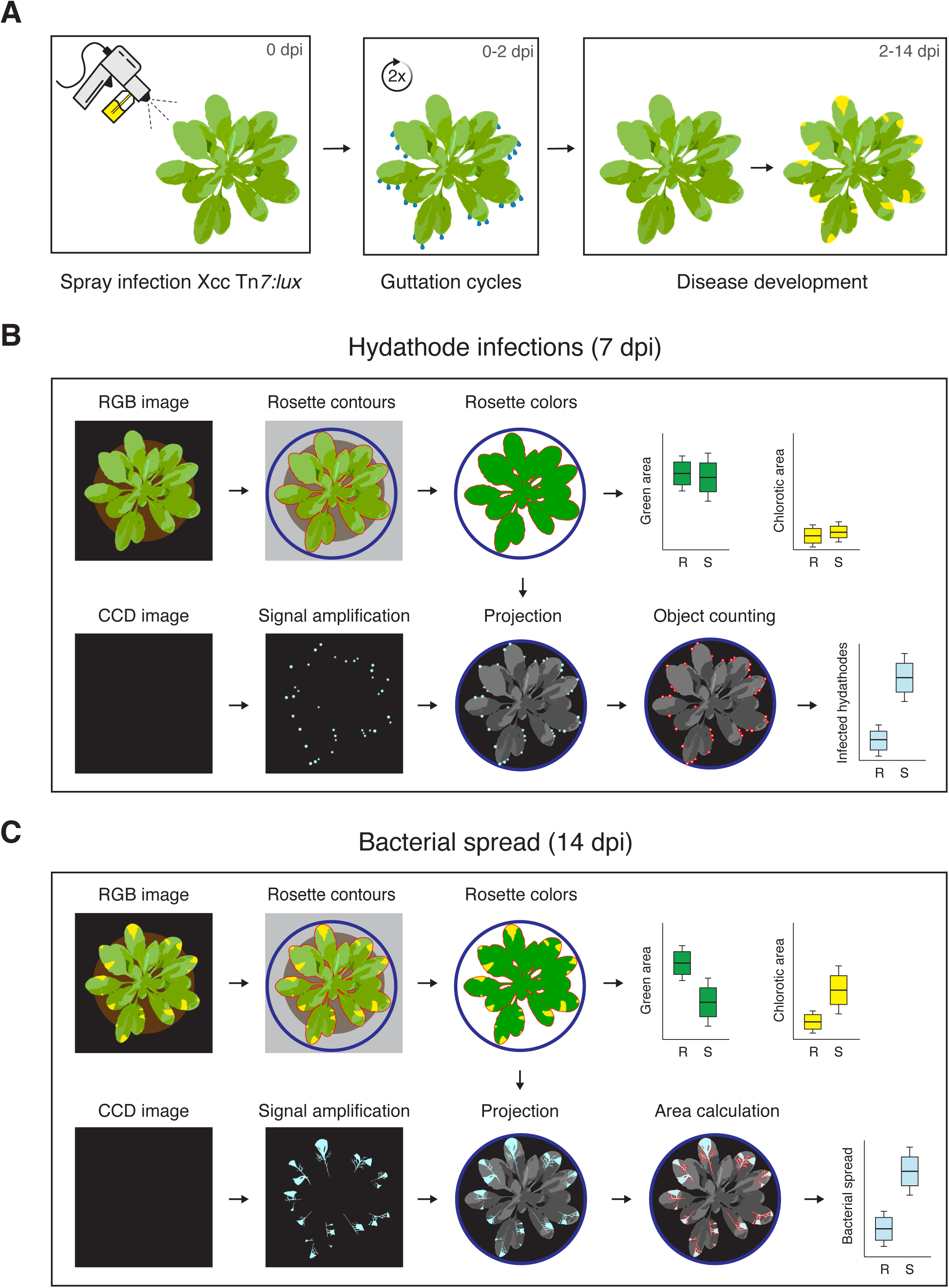
Schematic representation of the spray inoculation procedure with Xcc Tn*7:lux* and the subsequent image analysis pipeline. **A)** Spray inoculation procedure with Xcc Tn*7:lux* mimics natural infection via hydathodes. **B)** Schematic overview of the pipeline for quantification of infected hydathodes per rosette at 7 dpi resulting in number of infected hydathodes, total rosette area and chlorotic area as outputs. R, resistant; S, susceptible. **C)** Schematic overview of pipeline for quantification of disease symptoms and bacterial spread per rosette at 14 dpi resulting in bacterial spread (defined as total luminescent area), total rosette area and chlorotic area as outputs.

### Automated image analysis pipeline: developing the software

To automatically collect quantitative data, an image analysis pipeline (Figure 1B and C) was written in Python using the OpenCV Library (Bradski and Kaehler, 2000). To start the analysis, the output images from the two cameras are calibrated using a 4×11 asymmetric dots calibration sheet allowing us to overlay both images. In the RGB image, individual rosettes are then detected, outlined and sorted based on user-adjustable HSV color values and contouring parameters. In addition, the total rosette size and the number of green and yellow pixels are quantified. Next, using the defined rosette contours, the smallest enclosing circle is used as a mask around each rosette for the CCD image. Within each circle, luminescence is quantified by the number of luminescent objects and the luminescent area in pixels. To eliminate noise and artefacts in the CCD image, a luminescent signal is only considered when the pixels fall within a user-defined size range (e.g. minimum two pixels). Finally, all quantified variables for each rosette are saved in a .csv file and an overlay .png image of the RGB and CCD image is created for visual inspection. The pipeline features an environment file in which the different parameters can be adjusted to optimize the pipeline for other setups. All available parameters, code and instructions for this pipeline are provided on GitHub (https://github.com/MolPlantPathology/Digital_phenotyper).

### Digital phenotyping quantifies disease severity at different stages of infection

To confirm the validity of our method, we benchmarked our digital phenotyping pipeline against other well-established methods. To do so, we performed spray inoculations of Xcc8004 Δ*xopAC* Tn*7:lux* on three Arabidopsis genotypes that we showed previously to have different levels of disease susceptibility; two natural accessions Col-0 (resistant) and Oy-0 (hypersusceptible) and Col-0 mutant line *sut1-9*, which carries a knockout mutation in the CNL-type plant immune receptor SUT1 and shows an intermediate disease resistance phenotype against Xcc8004 Δ*xopAC* (Paauw et al., 2023; Taks et al., 2024). As the three different levels of resistance against Xcc can already be observed for these genotypes at the first stage of the infection in the hydathodes, it allows us to benchmark the sensitivity of our digital phenotyping setup to detect and quantify immune responses in hydathodes. Using our established detached leaf assay (van Hulten et al., 2019), we were able to detect that both *sut1-9* and Oy-0 display an increased number of hydathode infections at 7 dpi based on the number of luminescent hydathodes observed per leaf (Figure 2A). In parallel, the number of hydathode infections was quantified per rosette using the new pipeline (Figure 2B). Both methods yielded significant differences in hydathode colonization by Xcc between the three Arabidopsis lines. Next, to quantify the level of resistance at later stages of infection (i.e. in the vasculature and mesophyll), the proportion of the leaf area colonized by Xcc was quantified in the detached leaf assay using the ScAnalyzer script (Figure 2C) (Paauw et al., 2024) and with our digital phenotyping setup imaging the whole rosette at 14 dpi (Figure 2D). Again, at 14 dpi, we noted a consistent correlation between the results obtained with the earlier used detached leaf assay and the non-invasive imaging of whole rosettes. A significant difference between the genotypes was observed with a smaller sample size (Figure 2B and D, n = 16; Supporting Information 1) compared to the detached leaf assay (Figure 2A and C, n = 63; Supporting Information 1) at both timepoints (7 and 14 dpi), indicating a high robustness of our approach. Besides bacterial colonization, we analyzed disease symptoms and compared whether both parameters give consistent results. Both quantification of chlorotic disease symptoms and bacterial colonization yielded significant differences between Col-0 and Oy-0 at 14 dpi, with bacterial colonization resulting in more robust data (Figure S2A), as bacterial spread precedes symptom development (Paauw et al., 2024)(Figure S2B). Therefore, utilizing bacterial colonization as phenotype can reduce the time between infection and phenotyping to obtain high-resolution data.

**Figure 2.**
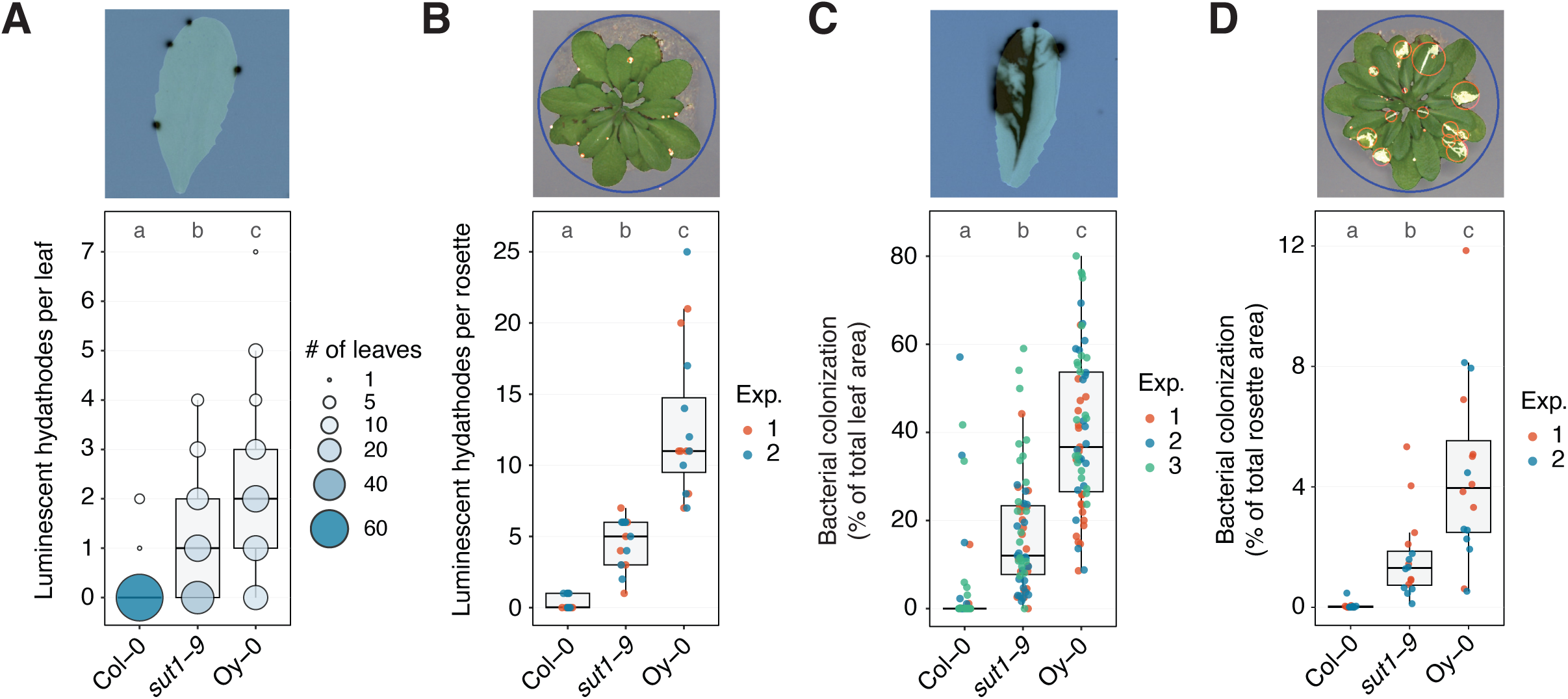
Digital phenotyping of entire plants is consistent with results of detached leaf assays using Xcc Tn*7:lux* infection of three Arabidopsis lines with varying disease susceptibility. **A)** Quantification of a *Xanthomonas* disease assay with detached leaves counting the number of hydathodes infected per leaf (n = 63 leaves per genotype). Leaves were imaged 7 days post spray inoculation (dpi) with Xcc8004 Δ*xopAC* Tn*7:lux.* Letters above the box indicate different significance groups based on a non-parametric Kruskal-Wallis test followed by Dunn’s Post-Hoc test, p-value threshold = 0.05. **B)** Same as panel A, except that whole rosettes were digitally imaged (n = 16 plants per genotype) and represent an overlay of RBG and luminescence picture (luminescence shown in false color). Each luminescent area is automatically highlighted by red circle. Letters above the box indicate different significance groups based on a non-parametric Kruskal-Wallis test followed by Dunn’s Post-Hoc test, p-value threshold = 0.05. **C)** Same as panel A, except that proportion of the leaf area colonized by *Xanthomonas* is shown per detached leaf 14 dpi (n = 63 leaves per genotype). Significance letters from non-parametric Kruskal-Wallis test with Dunn’s Post-Hoc test, p-value threshold = 0.05. **D)** Same as panel A, except that relative rosette area colonized by *Xanthomonas* is shown using digital imaging of whole rosettes (n = 16 plants per genotype). Letters indicate significance groups based on a non-parametric Kruskal-Wallis test followed by Dunn’s Post-Hoc test, p-value threshold = 0.05. Colored dots indicate data points from independent experiments.

While quantification at two defined timepoints yields robust data to assess the level of disease resistance, time-series measurements can capture the bacterial progression in the plant tissues with even more precision (Video S1). Moreover, in the case of Xcc infection in Arabidopsis, by imaging the infections over time, it is possible to study the dynamics of the infection process of individual hydathodes, yielding true single hydathode data (Figure 3). Together, we conclude that this digital phenotyping setup provides consistent data similar to the detached leaf assay using light-sensitive films (Paauw et al., 2024) at different timepoints.

**Figure 3.**
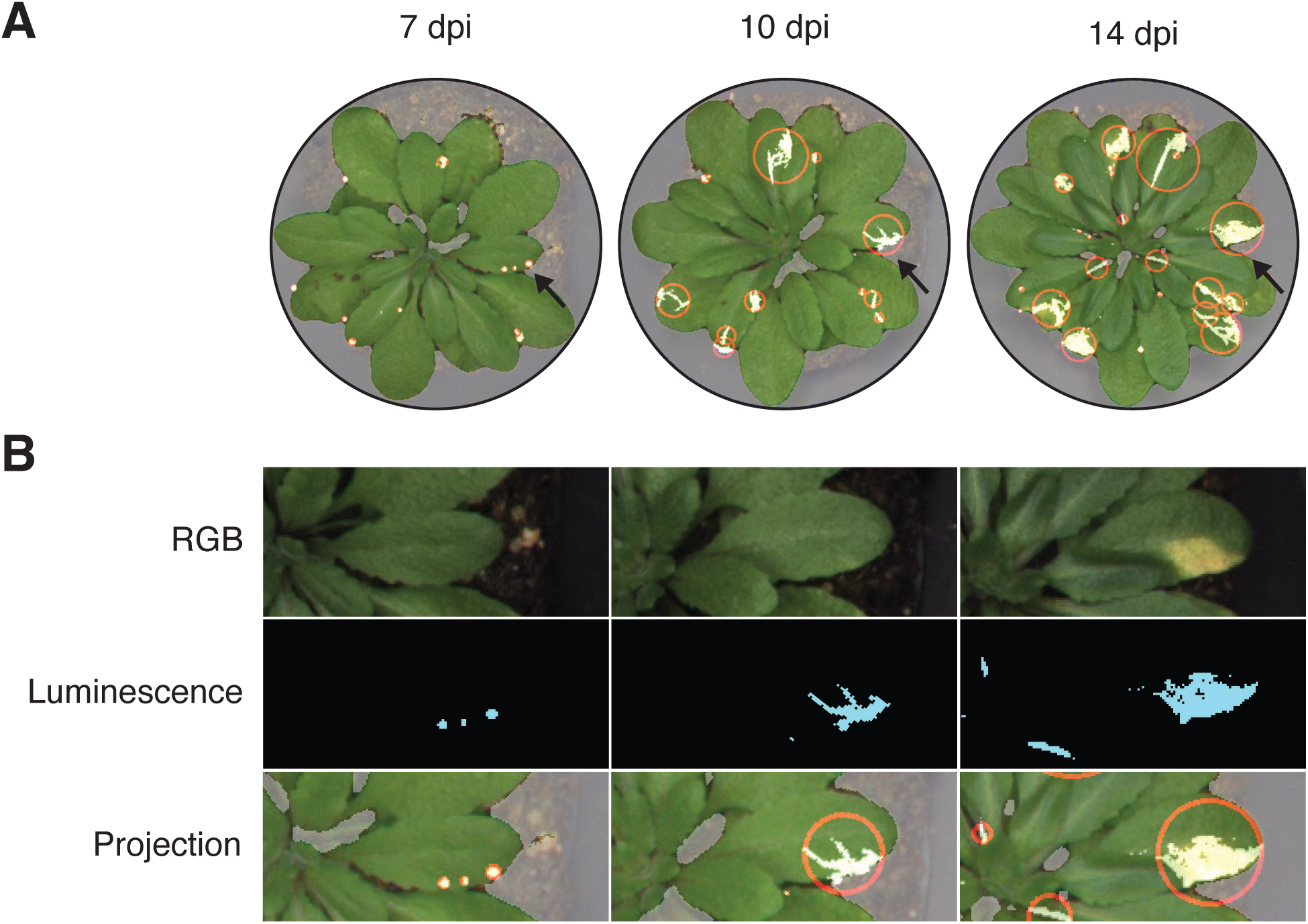
Non-invasive phenotyping allows for tracking of *Xanthomonas* outbreaks from a single hydathode. **A)** Imaging of Xcc Tn*7:lux* infections on *Arabidopsis* imaged at 7, 10 and 14 dpi in the digital phenotyper. Luminescence is shown as false color. Black arrow marks a single leaf enlarged in panel B. Individual luminescent areas are automatically marked with red circle. **B)** Enlarged images of leaf indicated in panel A showing Xcc outbreak from three individual hydathode infections in a single leaf.

### Digital phenotyping approach is flexible and applicable to mesophyll pathogens

Although our pipeline was optimized for the interaction between Xcc and Arabidopsis, we assessed its versatility using the model organism *Pseudomonas syringae* pv. tomato (Pst) strain DC3000 (PstDC3000) to look at stomatal infections in Arabidopsis (Melotto et al., 2017). Arabidopsis Col-0 is susceptible to PstDC3000 (Velásquez et al., 2017). To infer the ability of our pipeline to quantify resistance to Pst, a heterologous effector delivery system (EDV) was used (Fabro et al., 2011). In this approach, Pst was used to secrete the Xcc type III effector (T3E) XopAC, as it triggers an immune response in Col-0 accession. This response restricts bacterial proliferation through recognition of XopAC by the RKS1-PBL2- ZAR1 resistosome (Guy et al., 2013; Wang et al., 2015, 2019). Similar to Xcc, Arabidopsis plants can be spray-inoculated using PstDC3000 Tn*7:lux* and the luminescent signal can be imaged (Figure 4A). To quantify bacterial proliferation of Pst, we sprayed Col-0 wildtype plants and *rks1* and *zar1* knockout mutants with non-luminescent Pst, either wildtype or expressing *xopAC*, and determined the number of colony-forming units (CFU) in leaf discs of inoculated rosettes (Figure 4B). In wildtype Arabidopsis (Col-0), Pst expressing *xopAC* showed reduced bacterial proliferation at 3 dpi compared to the wildtype Pst. This reduced bacterial proliferation was not detected in the *zar1* and *rks1* knockout lines, indicating that host delivery of XopAC by the EDV system successfully triggers a ZAR1/RKS1-dependent immune response in wildtype Arabidopsis Col-0 plants. Next, to compare these results to the new image analysis pipeline, we sprayed the same Arabidopsis lines with Pst Tn*7:lux* (carrying either pEDV:*XopAC* or not) and quantified luminescence of whole rosettes (Figure 4C). The total bioluminescent area of Pst expressing x*opAC* was reduced in wildtype Col-0 plants but not in *zar1* and *rks1* knockout lines, suggesting that our digital phenotyping method consistently replicated the results of CFU counting. Therefore, we conclude that the phenotyping setup can also be applied to infer bacterial titers of mesophyll pathogens, highlighting the flexibility of the system.

**Figure 4.**
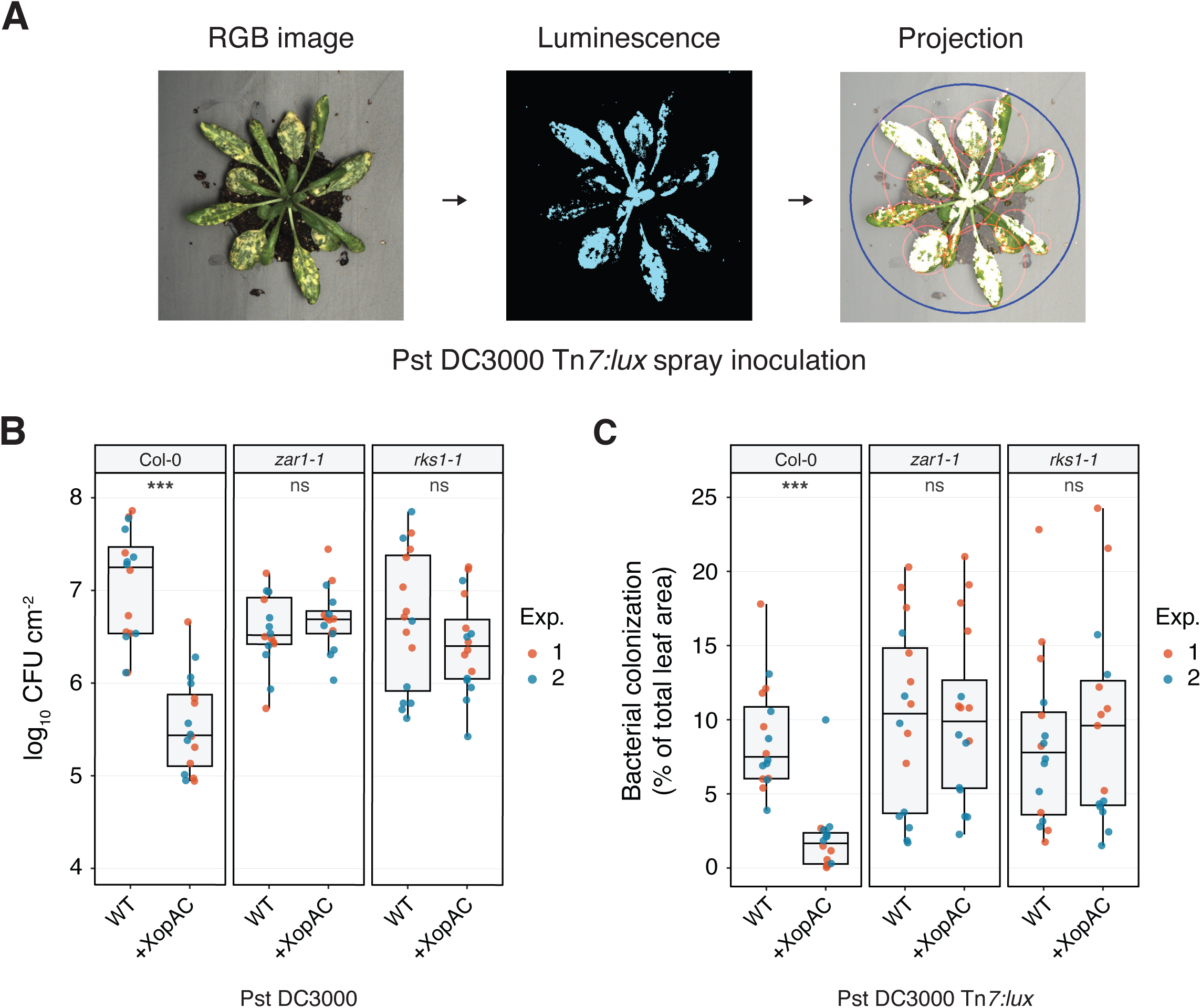
Quantification of disease susceptibility for the mesophyll pathogen *Pseudomonas syringae* using the digital phenotyping pipeline. **A)** Exemplary images taken with the digital phenotyper and output of the image analysis pipeline. Images were taken 5 days post spray inoculation (5 dpi) with Pst DC3000 Tn*7:lux* on Arabidopsis. **B)** Disease severity measured as bacterial counts, i.e. log_10_ CFU counts per cm^2^ leaf at 3 days following spray inoculation with wildtype Pst DC3000 or carrying pEDV6:*xopAC* on wildtype Arabidopsis (Col-0) and the loss-of-function mutants *zar1* and *rks1* (n = 16 plants per treatment). XopAC-recognition in Col-0 leads to reduced bacterial titers when Pst expresses *xopAC*, while this immune recognition is compromised in *zar1* and *rks1*. Statistics on comparison between WT and +XopAC result from a Student’s T-test, p- value *** <= 0.001, ns >= 0.05. **C)** Bacterial colonization of whole rosettes at 5 days following spray inoculation with Pst DC3000 Tn*7:lux* (carrying pEDV6:*XopAC* or not) on *Arabidopsis* Col-0 wildtype, *zar1* and *rks1* knockout lines (n = 16 rosettes per treatment). Treatments and results are consistent with phenotyping method shown in A). Significance on comparison between WT and +XopAC result from a non-parametric Wilcoxon rank test, p-value *** <= 0.001, ns >= 0.05.

## Discussion

Here, we present a non-invasive digital phenotyping method for quantification of plant disease symptoms and bacterial spread using bioluminescent reporter strains. Previous efforts, including from our lab, have shed light on early hydathode infections using bioluminescent reporter strains (Cerutti et al., 2017; Paauw et al., 2023; Taks et al., 2024). We recently developed an automated image analysis pipeline, called ScAnalyzer, for analyzing bacterial spread using detached leaves (Paauw et al., 2024). The approach presented here further improves this work by providing a method for non-invasive, highly automated phenotyping that yields accurate data on bacterial spread inside intact Arabidopsis rosettes (rather than only a single leaf). This digital phenotyping approach produces reproducible data that aligns with results obtained through conventional methods and is applicable to phytopathogenic bacteria that exhibit different infection strategies.

As shown in this work, our phenotyping setup can distinguish between several degrees of resistance against Xcc in Arabidopsis. This is important as many plant resistance mechanisms are quantitative in nature and polygenic (Corwin and Kliebenstein, 2017). In addition, plant pathogens are known to evolve rapidly, resulting in the repeated emergence of resistance breaking strains (de Wit, 2016; Di et al., 2017; Lenman et al., 2016). Therefore, being able to better screen for partial or quantitative resistance could provide novel genetic sources to stack different resistance genes against plant pathogens. In addition to acquiring highly accurate phenotyping data for both the plant and the pathogen, the setup is also suited to screen large Arabidopsis populations. As 16 plants can be imaged within five minutes, this allows for non-invasive screening of hundreds of plants in a single day. The image analysis pipeline is adjustable to other tray formats, allowing screening up to 40 plants at once when younger Arabidopsis plants are imaged, increasing the throughput even further. In this way, we can tap into the large resource of natural variation that is known to exist for Arabidopsis that can be exploited to screen for resistance (Weigel and Mott, 2009). Many previously reported methods for the detection of bioluminescent pathogens in plants are limited to assessing several leaves or plants at a time (Mutka et al., 2016; Wang et al., 2024; Xu et al., 2022). Furthermore, besides placing plants in the imaging device, the imaging setup and subsequent analysis pipeline both require little to no additional work, paving the way for implementation in a conveyer belt-like plant-pathogen-phenotyping platform.

In a complex multi-stage interaction between a plant host and its pathogen, such as Xcc infecting Arabidopsis, gaining knowledge on the different stages (i.e. hydathode, vasculature, mesophyll) provides a more holistic view on the entire disease cycle. For example, plant immune responses against Xcc occur as early as the hydathode colonization (Cerutti et al., 2017; Paauw et al., 2023; Taks et al., 2024), but also in the vasculature (Lin et al., 2022; Xie et al., 2023) and mesophyll tissue (Wang et al., 2015). The here described phenotyping platform is able to record plant resistance at individual stages of the Xcc infection or to perform e.g. a genome-wide association studies (GWAS) screening for candidate susceptibility genes. Moreover, the option of tracking a single infection event over time, provides the possibility to investigate the mechanism underpinning bacterial escape from the hydathode towards the xylem.

While this setup was designed to study the highly specialized vascular plant pathogen Xcc, we also show its versatility by unveiling resistance response against the model plant pathogen Pst infecting the leaf mesophyll (Xin et al., 2018). Using the same parameters as for the quantification of Xcc in later stages of infection (i.e. the mesophyll stage), we accurately recapitulate the known differences in resistance levels between several well-characterized Arabidopsis mutant lines. Furthermore, by comparing our method to the more traditional colony counting method, we found a correlation between the bioluminescence data and bacterial titers. The adaptability of this system provides ample opportunity for other pathogens that can be genetically tagged with bioluminescence, such as other *Pseudomonas* species and *Ralstonia solanacearum* (Jutras et al., 2021; Mutka et al., 2016; Xu et al., 2022). Therefore, with this setup, we expect to gain more detailed knowledge on the complex infection strategies used by bacterial pathogens of Arabidopsis.

### Experimental procedures

#### Plant growth conditions

All *Arabidopsis thaliana* lines used are listed in Supporting Information 2. For all experiments, seeds were stratified for 3-4 days on wet filter paper at 4 °C in the dark, and then sown in 40-pot trays (Desh Plantpak, Tray Danish size 40 cell (8×5)) in slitpots (Pöppelmann Teku, S 5,5 LB), using potting soil (Jongkind Substrates, Hol80 zaaigrond Nr 1) to which 500 ml Entonem suspension (Koppert Biological Systems, ± 1.6 x 10^6^ third stage *Steinernema feltiae* nematode larvae) was added per tray. Hereafter, trays were placed in short day conditions (11/13h light/dark, 22 °C, RH 70%). For the first 4-5 days, the trays were covered with a transparent plastic dome to ensure uniform germination.

#### Plasmid and transgenic strain generation

All bacterial strains, plasmids and oligonucleotides used are listed in Supporting Information 2. Bacterial strains were tagged with a double bioluminescence/fluorescence cassette using the mini-Tn7 transposon system (Choi and Schweizer, 2006; Soldan et al., 2021). Recipient strains (Xcc8004 Δ*xopAC* or Pst DC3000) were co-incubated on LB plates with one donor *E. coli* strain (DH5α + pRS-Tn7-pNPTII::*lux-pA1::mTq2* (Paauw et al., 2023; Soldan et al., 2021)) and two helper *E. coli* strains; one carrying Tn7 transposon genes *tnsABCDE* (DH5α + pUX-BF13 (Bao et al., 1991)) and one carrying a plasmid that facilitates conjugation (HB101 + pRK2073 (Figurski and Helinski, 1979)). Next, cultures were collected and transformants were selected on LB plates with antibiotics (10 μg/ml gentamicin and 50 μg/ml nitrofurantoin) and checked for luminescence using the ChemiDoc MP imager (Bio-Rad).

To create the pEDV6 (Fabro et al., 2011) vector carrying Xcc8004 *xopAC*, Xcc genomic DNA was isolated using the QIAGEN Puregene DNA-isolation kit for Gram-negative bacteria. Hereafter, the *xopAC* coding sequence was amplified using primers that contain *attB* sites for Gateway cloning. The amplified product was then cloned into pEDV6 by LR reaction creating pEDV6:*xopAC*. The resulting vector was transformed into PstDC3000 by electroporation (Choi et al., 2006).

#### *Xanthomonas* disease assays

Spray inoculations with bioluminescent Xcc8004 Δ*xopAC* Tn*7:lux-mTq2* (Xcc Tn*7:lux*) were performed as previously described (Paauw et al., 2023; van Hulten et al., 2019). Xcc strains were plated on KADO agar medium (Kado and Heskett, 1970) with antibiotics (10 μg/ml gentamicin and 50 μg/ml nitrofurantoin) and allowed to grow for 48-72 hours at 28 °C. Bacteria were collected, washed once with 10 mM MgSO_4_ and diluted to OD_600_ = 0.1. Silwet L-77 was added to a final concentration of 0.0002% and bacteria were sprayed on 4-to-5- week-old Arabidopsis plants (Preval sprayer, SKU# 0221). To promote natural infection, sprayed plants were subjected to two guttation cycles over two days in a climate chamber (Snijders Labs, MC1000) (van Hulten et al., 2019) and subsequently grown for an additional 12 days at short day light cycle (11/13h light/dark) at 24-22 °C with a relative humidity of 70% in the same chamber.

To quantify hydathode infections and bacterial colonization using light-sensitive film, at 7 and 14 dpi, the three most symptomatic leaves of each plant were taken and glued onto paper (A3 paper size) with a printed grid and covered with a transparent plastic sheet. Next, a light-sensitive X-ray film (Fuji Super RX) was placed on top and left for exposure overnight (van Hulten et al., 2019). The film was developed the next morning and scanned, alongside the leaves, in an A3 flatbed scanner (Epson Expression 12000XL). For 7 dpi samples, images were overlayed in Adobe Photoshop 2023/2024 to count luminescent hydathodes per leaf. At 14 dpi, scans of leaves and light-sensitive film were analyzed using the ScAnalyzer image analysis tool (Paauw et al., 2024) to quantify both the level of chlorosis and bacterial spread.

#### *Pseudomonas* disease assays

For both spray inoculations and infiltrations, Pst was grown overnight in 50 ml Falcon tubes containing 20 ml King’s B (KB) medium at 28 °C and 200 RPM. The next day, cultures were centrifuged, washed once with 10 mM MgSO_4_ and diluted to OD_600_ = 0.1 for spray inoculations or OD_600_ = 0.001 for infiltrations. For spray assays, Silwet L-77 was added to a final concentration of 0.002%. Next, 4-to-5-week-old *Arabidopsis* were sprayed (Preval sprayer, SKU# 0221) or pressure-infiltrated using a blunt syringe. Infected plants were grown for 5 days at SD conditions (11/13h light/dark, 24-22 °C, RH 70%). The first 3 days, plants were covered with a transparent plastic dome to promote infection.

To determine colony-forming units after leaf infiltration, at 3 dpi, two leaf discs (5 mm diameter) were taken from each infiltrated leaf with a leaf puncher, placed into 500 μl 10 mM MgSO_4_ containing two steel beads and homogenized twice for 1 minute at 30 Hz in a TissueLyser II (QIAGEN). Samples were serially diluted and plated in duplicates on KB agar plates containing 10 μg/ml gentamicin and 25 μg/ml rifampicin antibiotics. After 24h, colonies were counted and log colony-forming units per cm^2^ leaf material (log_10_ CFU cm^-2^) was calculated.

#### Disease quantification with digital phenotyper

At 7 and 14 dpi (Xcc) or 3 and 5 dpi (Pst), plants in slit pots were taken from their respective locations in the 40-pot tray and placed into an anodized (black) aluminium tray with 16 (4×4) positions for imaging (Figure S1B). Next, the plants were covered with a black plastic lid and kept in the dark for 5 minutes to reduce photosynthetic background signal when imaging luminescence. Hereafter, the imaging tray was placed in the digital phenotyper and was imaged in a standard program (300s exposure CCD camera and 70ms exposure RGB camera). Raw images (RGB and CCD) were exported and fed into our image analysis pipeline.

#### Automated image analysis

Raw images are processed by an in-house developed Python analysis pipeline that uses the open-source OpenCV Library for image analysis (Bradski and Kaehler, 2000). In this pipeline, rosettes are detected in the RGB image according to user-adjustable HSV values for leaf tissue. The user can define row and column numbers, indicating the number of rosettes to be found in the image. A sorting algorithm then orders the rosettes and assigns row and column numbers. Whenever less rosettes are detected than initially defined, the RGB image is brightened and saturated to increase detection capabilities of the algorithm. Next, each rosette contour is drawn and masked to select only plant tissues and no background (e.g. soil or tray). For each masked rosette, several output parameters are calculated, including rosette area, green area and yellow area based on user-defined color parameters. A smallest-enclosing circle is then drawn around each rosette mask to create a mask for the luminescence image. The luminescence image is overlayed onto the RGB image using the homography matrix, aligning both RGB and CCD cameras, and for each smallest-enclosing circle, the number of luminescent objects (corresponding to colonized hydathodes (Xcc: 7 DPI)) and/or total luminescence signal (Pst: 3 and 5 DPI, Xcc: 14 DPI) is calculated. Metadata on each rosette (plant genotype, treatment etc.) can then be matched to all measurement values and combined into a data output file. All code and instructions on parameter selection and how to perform the image analysis are freely available on GitHub (https://github.com/MolPlantPathology/Digital_phenotyper).

#### Quantification and statistical analyses

All data analyses and plotting were done in R (v.4.3.2). For statistical analysis of parametric data, a two-way ANOVA followed by a Tukey post-hoc test was performed for multiple comparisons between groups and a Student’s T-test when comparing two groups. For non-parametric data, such as hydathode counts or proportional data, a Kruskal-Wallis test followed by a Dunn’s post-hoc test was performed for multiple comparisons and a Wilcoxon test when comparing two groups. All experiments were performed in at least two independent repetitions with similar results. Data from repetitions was pooled for visualization and statistical analysis.

## Acknowledgements

The authors would like to thank Ludek Tikovsky and Harold Lemereis for excellent plant care in the greenhouse. Also, we would like to thank Dominique Roby and Darrell Desveaux for sharing Arabidopsis mutant lines and Alan Collmer, Gail Preston and Laurent Noël for sharing bacterial strains and plasmids. Lastly, we would like to thank LemnaTec for the assistance in designing the digital phenotyper. This research was supported by the Topsector T&U TKI programs “Stop natural entry” and “Finding the Achilles’ heel of Brassica for Black Rot disease” (grants EZ-2012-02 and TU18024 to H.A.v.d.B.) and the breeding companies Bejo Zaden B.V. and Rijk Zwaan.

## Data availability statement

The data that support the findings of this study are available from the corresponding author upon reasonable request.

## Supporting Information legends

**Figure S1. Hardware used for the imaging of Arabidopsis rosettes and bacterial luminescence**

**A)** Digital phenotyping cabinet with dual RGB/CCD camera system. **B)** 3D model used for production of aluminum 16-pot imaging trays. Distances depicted in mm. **C)** Schematic overview of 16-pot imaging tray layout and output of both RGB and CCD cameras. Bacterial luminescence in CCD projection depicted in cyan.3D model used for production of aluminum 16-pot imaging trays. Dimensions depicted in mm.

**Figure S2. Bacterial colonization precedes chlorotic disease symptoms and shows more pronounced differences between Col-0 and Oy-0 accessions when acquired at 14 days post inoculation.**

**A)** Quantification of a *Xanthomonas* disease assay using digital imaging of whole rosettes by either (*left*) chlorotic disease symptoms or (*right*) bacterial colonization (luminescence). Rosettes (n = 32 per genotype) were imaged 14 days post spray inoculation (dpi) with Xcc8004 Δ*xopAC* Tn*7:lux.* Letters above the box indicate different significance groups based on non-parametric Wilcoxon test. Differences between Col-0 and Oy-0 accessions are more pronounced when analyzed by bacterial colonization rather than chlorotic disease symptoms. **B)** Representative rosettes of the quantification shown in panel A). *Top*, RGB images of whole rosettes showing chlorotic disease symptoms; *bottom*, false coloring of luminescent signal in the same rosettes. Individual luminescent areas are encircled. The presented data suggests that bacterial colonization precedes chlorotic disease symptoms.

**Video S1. Time series of Xcc Tn*7:lux* infection in susceptible Oy-0 from 5 to 19 days after inoculation**

The Oy-0 rosette was spray-inoculated with Xcc Tn*7:lux* and then imaged twice per day from 5 to 19 dpi (luminescent signal in orange). Separate images were joined into an animated GIF image.

**Supporting Information 1. Exact p-values between groups calculated by non-parametric Kruskal-Wallis test for all panels in Figure 4**

**Supporting Information 2. All plant lines, bacterial strains and recombinant DNA used in this study**

