## Supplementary figures and images for "Visualization and quantification of spatiotemporal disease progression in Arabidopsis using a bioluminescence-based imaging system"

### Supplemental Figure S1

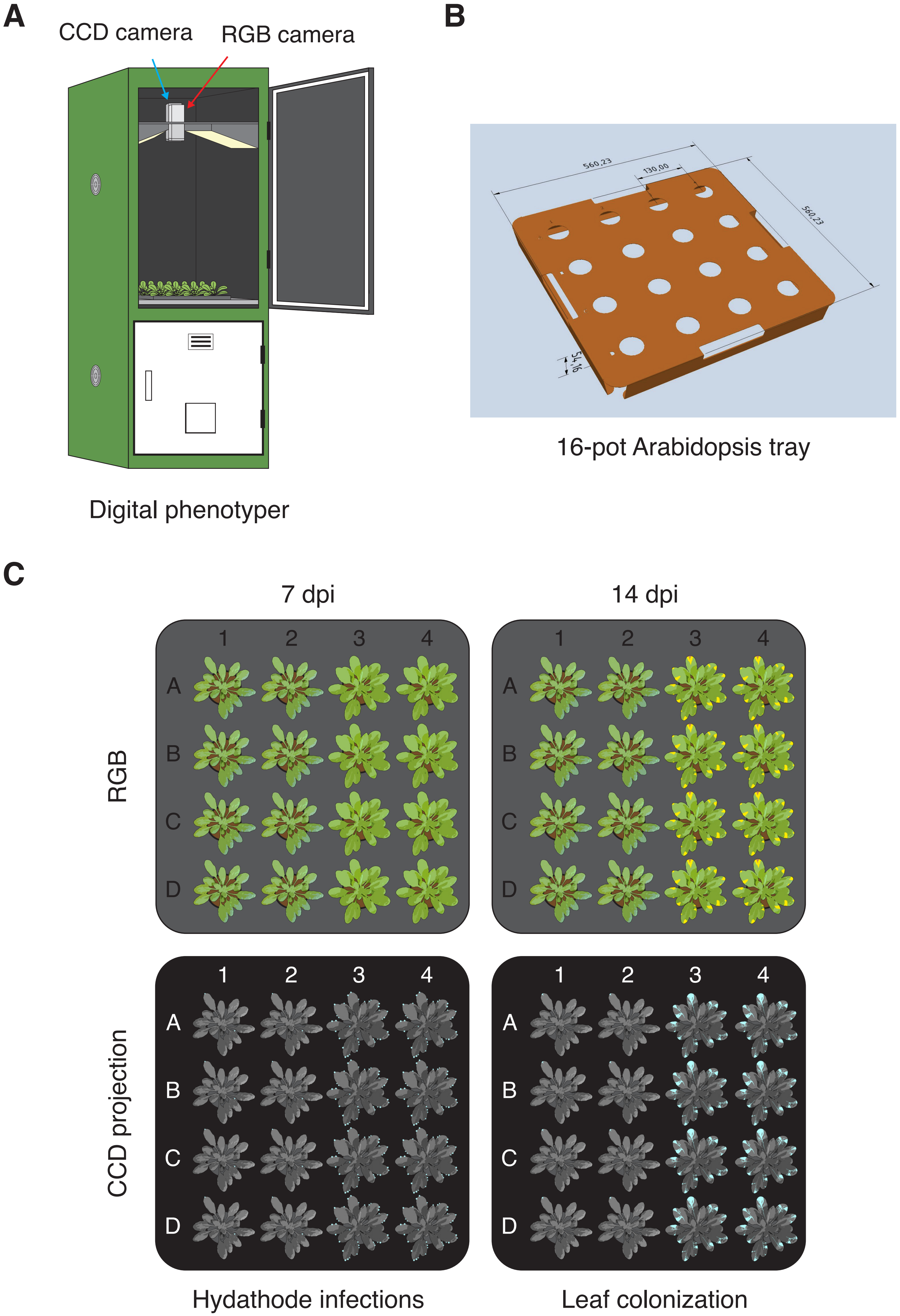

### Supplemental Figure S2

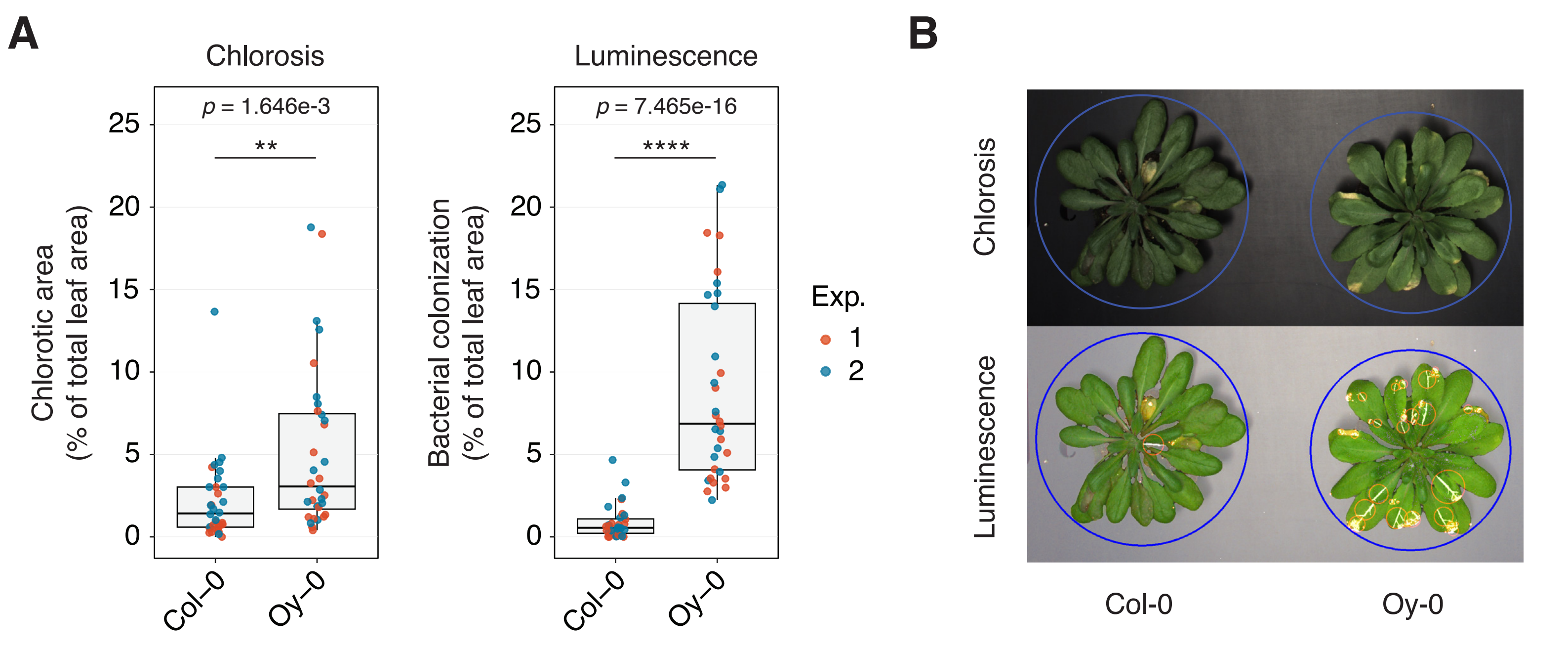

### Supplemental Video S1

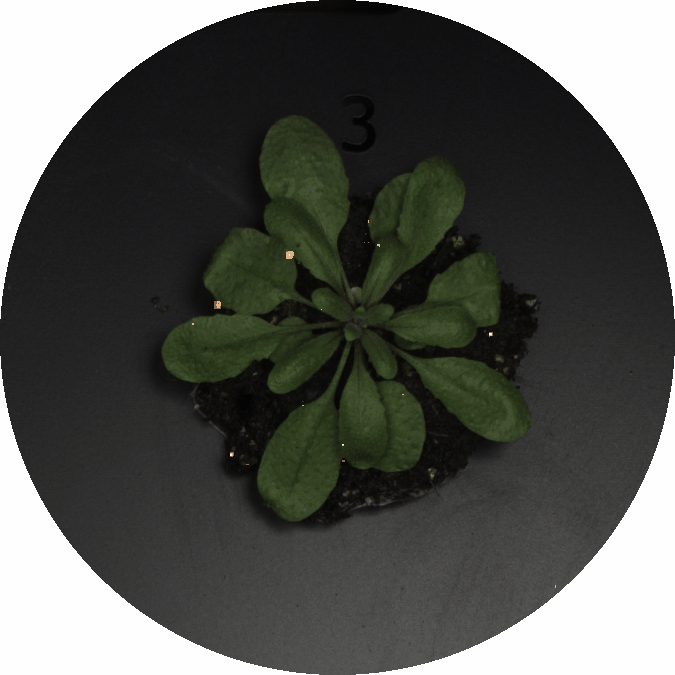
